# Modelling habitat suitability for multiple priority weed species to predict invasion hotspots for strategic management in complex landscapes

**DOI:** 10.64898/2026.05.31.729134

**Authors:** Kyana N Pike, Wen-Hsi Yang, Ben Gooden, Luke O’Loughlin, Richard Milner, Brett Howland, Steve Taylor, Jens Froese

## Abstract

Established invasive alien plant species require ongoing, costly management to reduce harm to agricultural and environmental values. Heterogeneous landscapes are often under threat from multiple long-established invasive plants, whose simultaneous management presents strategic and tactical challenges. Systematic monitoring of weed populations enables more strategic management by providing more detailed insights into invasion threats than the more readily available presence-only data. We leveraged six years of weed density monitoring and high-resolution environmental data to model habitat suitability for four high-priority weeds across the Australian Capital Territory. We developed habitat suitability maps for African lovegrass, Chilean needle grass, serrated tussock grass and St John’s wort. Maps were then standardised and combined to derive potential multi-species invasion hotspots at different management thresholds and explore patterns of secondary invasion threat. While some species shared environmental drivers such as elevation, each responded to distinct combinations and interactions among environmental drivers. Overall, we found approximately 71% of land had suitability for multiple weeds at various densities, but only 11% is suitable for two or more species to reach high densities. Areas suitable for most weeds often had St John’s wort as the next most suitable invader. These findings could improve management efficiency and reduce intervention challenges by focussing integrated control across species in a small number of priority areas. These findings support spatially explicit, multi-species decision-support tools for surveillance, prioritisation and strategic landscape-scale control.

## Introduction

Invasive alien plants (hereafter referred to as ‘weeds’) are a major threat to biodiversity, ecosystem health and agricultural productivity worldwide (Vilà et al. 2011; Pyšek et al. 2012; IPBES 2019, 2023). In many regions, weeds have become so widespread and dominant in their introduced range that they require ongoing management in perpetuity, often at considerable cost, to limit harm to agricultural and environmental values. In Australia for instance, widespread weeds in rangeland systems cost an estimated AUD 3.9 billion per year in lost production and control activities (Humphries and Florentine 2021). Despite this, the sustained management of long-established, widespread weeds remains a challenge in existing science and management frameworks, which historically focused on new incursions and factors influencing establishment and spread (Catford et al. 2011). Although substantial advances have been made in understanding invasion dynamics, this knowledge does not always translate into information that is directly usable for on-ground management of established invasions (Kuebbing et al. 2013).

From local to landscape scales, weeds interact with their environments in complex and spatially variable ways, making management across heterogenous landscapes particularly challenging. These spatial dynamics complicate decisions about where to intervene within established populations to reduce ongoing invasion pressure, and where to prioritise surveillance to detect new incursions. The threat of an invasive weed may vary, as not all areas that can support establishment will sustain high-density infestations, meaning the impacts of invasive weeds can vary markedly across landscapes (O’Neill et al. 2021). When time and resources for control are limited, effective management of widespread species therefore depends on prioritising interventions that deliver the greatest benefit. This in turn requires an understanding of how invasion threat varies spatially, allowing management to focus on areas where intervention is both most urgent and most likely to support recovery.

Adding to the management challenge is that virtually all ecosystems contain multiple weeds, meaning an increased likelihood of multiple long-established species requiring on-going management (Kuebbing et al. 2013; Pearson et al. 2016a, b). How weeds or their management may interact in a variety of positive or negative ways can make outcomes difficult to predict and limit the effectiveness of management (Lone et al. 2024). For instance, mutualism between weeds may promote high abundance of both (e.g. invasional meltdown (Simberloff and Von Holle 1999; O’Dowd et al. 2003), such that managing one species alone is unlikely to reduce overall abundance (O’Loughlin and Green 2017). Conversely, competitive interactions among invaders may supress high densities of one species, such that managing the dominant invader may release another to become highly invasive (known as management-mediated “secondary invasion”: Pearson et al. 2016b; Lone et al. 2024).

Secondary invasion is a very common outcome when managing invasive plants, with 75% of studies reporting an increase in secondary invader abundance after suppression of a dominant weed (Pearson et al. 2016b) rendering the efforts to reduce weed abundance ineffective.

Effective management for long-established and high-impact weeds requires multi-species frameworks, tools and approaches to better monitor and control for potential high-threat secondary invaders across regions of co-occurrence (Kuebbing et al. 2013; Pearson et al. 2016b; Lone et al. 2024).

Developing spatially explicit decision-support tools to help managers assess potential threats of co-occurring weeds requires an understanding of where priority species may overlap and reach high density at broad spatial scales. Habitat suitability models (HSMs) are a useful tool for predicting the potential spatial distribution of suitable habitats for target species within a given landscape (Guisan et al. 2017a). These models relate empirical observations of species presence or abundance to environmental predictors that can represent an ensemble of species’ niche requirements (Guisan et al. 2017a). HSMs have been applied extensively in invasive plant management to support decision-making across various environments, such as by predicting where new invasions could occur (e.g. Jarnevich et al. 2024), identifying broad environmental drivers of invasion (e.g. Catford et al. 2011) or forecasting future spread under climate change (e.g. Chai et al. 2016; Wang and Wan 2020; O’Neill et al. 2021; Beaury et al. 2023). The approach is also referred to by other terms, including species distribution models or ecological niche models, reflecting variations in methodology and application. A common limitation is the data upon which they are based. Most models rely on presence only data, supplemented with pseudo-absences, which limits their ability to provide insights into population demography or quantify potential severity of an invasion (Guisan et al. 2017b; Beaury et al. 2023). In contrast, models built with empirical density data are more informative for management, as they go beyond weed presence and can predict where species infestations may also be more severe in terms of higher density levels (O’Neill et al. 2021; Beaury et al. 2023). However, weed density data are labour-intensive to collect and therefore remain relatively scarce (O’Neill et al. 2021).

The overarching aim of this study was to provide spatially explicit insights for geographically relevant on-ground management decisions regarding the invasion risk posed by multiple co-occurring priority weed species. We leveraged a rich dataset to develop habitat suitability maps for four priority weed species, that allowed for assessments of both where weeds may grow and where is more suitable to support high-density infestations of these high threat weeds. The spatially explicit nature of these models also enabled additional down-stream analysis that can assist with more targeted and efficient multi-species management planning. Specifically, we (1) modelled habitat suitability for four priority, co-occurring weed species using spatial density data and boosted regression tree approaches, quantifying the potential suitability of the landscape to support low to high densities of each weed; (2) identified and mapped potential multi-species invasion hotspots, highlighting areas where environmental conditions may support multiple weeds, as well as areas where the threat of multi-species high-density invasions was greatest; (3) assessed patterns of secondary invasion threat within these multi-species hotspots by ranking species suitability scores where multiple species overlap to provide insights into which weed poses the greatest secondary invasion threat in relation to each potentially dominant weed. The aims were designed to support evidence-based, adaptive weed management strategies in heterogenous landscapes, enabling practitioners to prioritise weed surveillance and control based on predicted invasion risk across multiple co-occurring priority species. We developed this multi-species weed management framework using long-term weed population density data to support decision-tools in a management-relevant setting.

## Methods

### Study area

The Australian Capital Territory (ACT) is an ecologically diverse region in temperate southeastern Australia that is subject to continuous management for multiple high priority weeds. It encompasses low-lying plains in the northeast, heavily modified urban, peri-urban and agricultural landscapes, and higher elevation areas of largely intact vegetation in the Brindabella Ranges to the southwest. The urbanised landscape is characterised by low-density development interspersed with extensive green spaces and a network of conservation reserves containing remnant native vegetation (ACT Government 2018). These areas were historically dominated by natural temperate grasslands and box-gum grassy woodlands, now listed as threatened ecological communities. The Ngunnawal people as Traditional Custodians actively managed these landscapes for tens of thousands of years prior to European colonisation in the 1820s, which introduced many plant species that have since become invasive. Today, land managers are responsible for managing upwards of 150 weeds over the territory, with priority placed on those posing the greatest biosecurity risk, requiring sustained and coordinated management efforts across the region.

### Priority species

Here we focussed on four of the most pervasive weeds that threaten ecological communities and are well monitored: African lovegrass (*Eragrostis curvula*), Chilean needle grass (*Nassella neesiana*), serrated tussock grass (*Nassella trichotoma*) and St John’s Wort (*Hypericum perforatum*). These four species were introduced by the first half of the 1900s and are all established within the ACT. Due to their capacity to reach high densities and dominate ground cover, often forming monocultures, these four species are regularly controlled as priority species within the ACT. Because these species often co-occur, control of one can influence the invasion risk posed by others, complicating management decisions. Despite intensive efforts, these weeds persist and continue to threaten native ecosystems, highlighting the need for improved spatial planning and integrated control strategies.

### Invasive plant density data

We used a dataset containing density observations for the four priority species that was systematically collected from 1218 fixed monitoring points across the ACT. Over the course of a six-year monitoring period (January 2018 to March 2024) most points were revisited multiple times (median = 3 monitoring events per point range 1-7), producing a total of 4036 monitoring events in our dataset (see Fig. 1). Most monitoring events (94%) took place during spring to summer (October to February), with remaining monitoring distributed across other months when conditions allowed. For each monitoring point, the observer would measure the distance to the three closest individuals of a target weed within a 15m radius search area. Plant density per hectare was then estimated from these distance measures using a nearest-neighbour distance estimator (Pollard 1971; Laycock and Batcheler 1975; White et al. 2008). Upper and lower bounds were imposed on the density estimates to avoid unrealistic extremes: if less than three individuals of the target weed were present but at 15m or if three individuals were very close to zero. As a result, our density estimates were right censored at 9540.78 plants per hectare, and interval censored between 0 and 14.13 plants per hectare (see Supplementary Figure 1). At each monitoring event this process was repeated for each target invasive plant to give independent measures for all four invasive plants at the same location and time.

**Figure 1.**
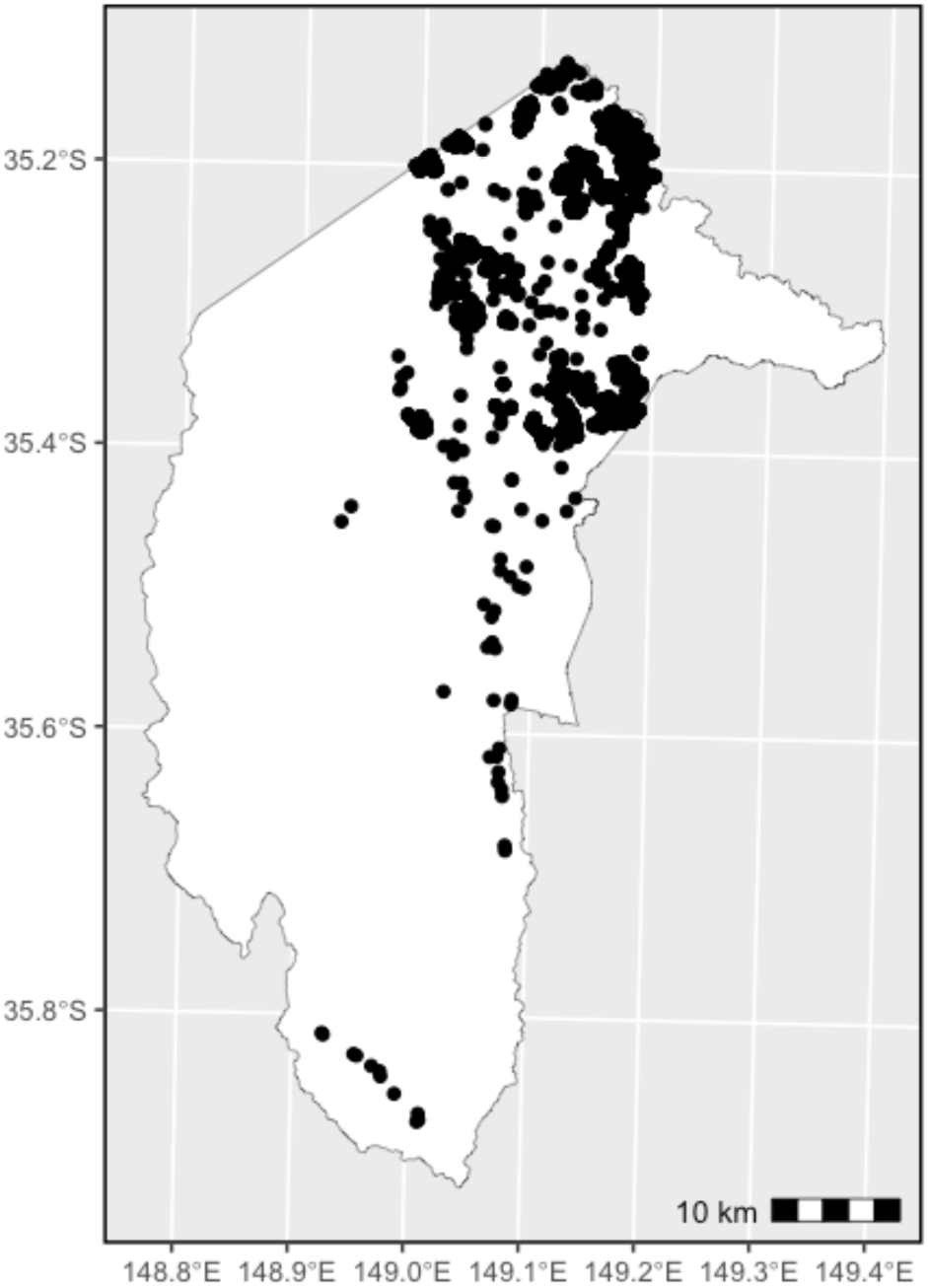
Locations of the 1218 fixed monitoring plots throughout the ACT where the density of African lovegrass, Chilean needle grass, St John’s wort, and serrated tussock grass was recorded over six years.

### Environmental covariates

We initially compiled a set of 57 variables known or suspected to influence the presence or density of at least one of the four priority weeds (see Supplementary Table 1). Some variables in this initial set exhibited strong multicollinearity (|r| = >0.90) or did not have appropriate geospatial data available at a resolution of at least 1km to enable prediction. To select appropriate variables, we then used a combination of data driven methods i.e. boosted regression trees from the ‘dismo’ package (Hijmans et al. 2023) and lasso regularisation using the ‘glmnet’ package (Friedman et al. 2010) and expert elicitation to confine this to 13 covariates for model inclusion. For example, water availability strongly influences plant growth, but capturing this effect at the appropriate temporal scale and metric is challenging when the relevant signal is not known *a priori*. To identify the most informative timespan and metric for rainfall patterns that impact weed density across landscapes, we compared total, mean, and standard deviation of rainfall over the 30, 90, and 365 days preceding monitoring. Although all nine variables were correlated, both model performance and expert assessment indicated that total yearly rainfall was the most informative.

The selected covariates represented topographic (elevation, % slope, topographic wetness index, easterness index, northerness index), climatic (total yearly rainfall, averaged over 2017 to 2023), soil (broad soil process, broad hydrogeological soil composition), land cover/use (vegetation type, projected canopy cover for trees >3m, land-use type) and disturbance factors (number of fires, distance to nearest road, track or trail) potentially promoting or limiting weed introduction, establishment and growth ( see Supplementary Fig. 2 and Supplementary Table 2 for details). Pairwise Pearson correlations (among continuous covariates) were all below |r| = 0.7 (maximum |r| = 0.63), indicating sufficiently low multicollinearity (Guisan et al. 2017c). All data were pre-processed to a common extent and resolution (1 hectare) using the ‘terra’ package (Hijmans 2025) in R version 4.4.1 (R Core Team 2024).

### Habitat suitability prediction

We modelled habitat suitability separately with boosted regression trees (BRTs) for each of the four priority weeds using the ‘gbm.step’ function from the ‘dismo’ package (Hijmans et al. 2023). BRTs are useful when modelling complex ecological data because of their ability to capture complex nonlinear and interactive covariate effects (Elith et al. 2008). However, default settings are rarely optimal for real ecological datasets and often require calibration to avoid overfitting (Chollet Ramampiandra et al. 2023).

For each priority species we held all 13 covariates constant and tested 144 combinations of three key hyperparameters: 1. The learning rate, or shrinkage parameter which determines the contribution of each tree to the growing model; 2. Tree complexity which determines the number of interactions that are permitted and 3. The bagging fraction, or the proportion of training data selected at each iteration (Elith et al. 2008). We selected parameter values based on the recommendations of Elith et al. (2008) and included nine tree complexity values (1 through to 9) along with four bagging fractions (0.6, 0.65, 0.7, 0.75), and four learning rates (0.001, 0.005, 0.01, 0.03). All models were fit with a Gaussian error structure for the natural logarithm of weed density, and a maximum of 25,000 trees with 10-fold cross validation.

For each species, we then selected the parameter combination that best balanced improvements in predictive performance against overfitting (Elith et al. 2008; Elith and Leathwick 2017; Chollet Ramampiandra et al. 2023; Yates et al. 2023). We first discarded any models with less than 1000 trees (Elith et al. 2008). Our wide range of tree complexity values (1 – 9) enabled us to identify at which two tree complexity values the out-of-sample root mean squared error (RMSE) values stabilised (see Supplementary Figure 3). We then reviewed a subset of models with these two tree-complexity values, and selected the best one based on the smallest out-of-sample RMSE and minimum % difference with its in-sample RMSE. The selected models were then used to map predicted potential weed densities per hectare across the ACT using the ‘terra’ package v 1.8-54 (Hijmans 2025). To provide an index of suitability that supports comparison of relative suitability across the four priority weed species, we rescaled each species’ density map using min-max normalisation to produce a habitat suitability index (HSI) ranging from 0 to 1. The model-predicted index can be interpreted as indicating the suitability of a given habitat in supporting relatively higher or lower population densities of the target species, where 0 indicates the environment is not suitable for the weed at a one hectare resolution.

### Hotspots for multi-species invasions

Next, we defined two threat thresholds from the distribution of rescaled HSI values for each species and then identified areas where multiple species overlap at each threshold. To determine threat thresholds we first, removed cells of the predicted map deemed unsuitable (i.e. areas where HSI < 0, which is where plants per ha/cell<1 ). Second, we calculated the 1^st^ quartile (Q1) and 3^rd^ quartile (Q3) of the remaining HSI values (see Supplementary Figure 4 for details). Third, we identified locations where two or more priority species exceeded the Q1 (i.e. suitability scores where only the lowest quarter of values were discarded to include a mix of low to high suitability) and Q3 (i.e. suitability scores where only the highest quarter were retained to include high values only) thresholds respectively. These locations can be interpreted as representing hotspots for multi-species invasion to management-relevant areas where the weed may be present at variable densities versus only where it may reach more severe densities respectively. This is more likely to represent realistic scenarios where managers have resourcing to target weeds at mixed densities except where weeds are at their lowest (>Q1), versus when resourcing or other constraints lead to prioritising only the highest density (>Q3) areas.

Fourth, for all identified multi-species invasion hotspot locations we ranked the four priority species according to their relative HSI values. Rankings ranged from 1 (highest HSI) to 4 (lowest HSI). To test whether species differed in how often they ranked highest in predicted suitability across the ACT, we applied a chi-squared test to the frequency distribution of rank-1 outcomes using the ‘stats’ package (R Core Team 2024). We then assessed whether the species most frequently ranked first exhibited similar suitability indices to the species most frequently ranked second by calculating Spearman’s rank correlation between their HSI values across all locations that met either threshold (base R; R Core Team 2024).

## Results

### Model tuning

Overall, we observed moderate to high R^2^ for our candidate models (SJW = 0.49, ALG = 0.62, ST = 0.45 and CNG = 0.67), suggesting that the selected covariates were generally informative for characterising invasive plant density across monitoring plots. However, these findings also suggest there remains unexplained variation that the 13 environmental covariates did not capture. Furthermore, the best selected models had tree complexities between 3 – 4 indicating that complex interactions between environmental covariates are likely and important to consider (Table 1). Predictive performance varied among species (Table 1), however, across all species, the models captured general spatial patterns of relative suitability despite uncertainty at the extremes.

**Table 1.**
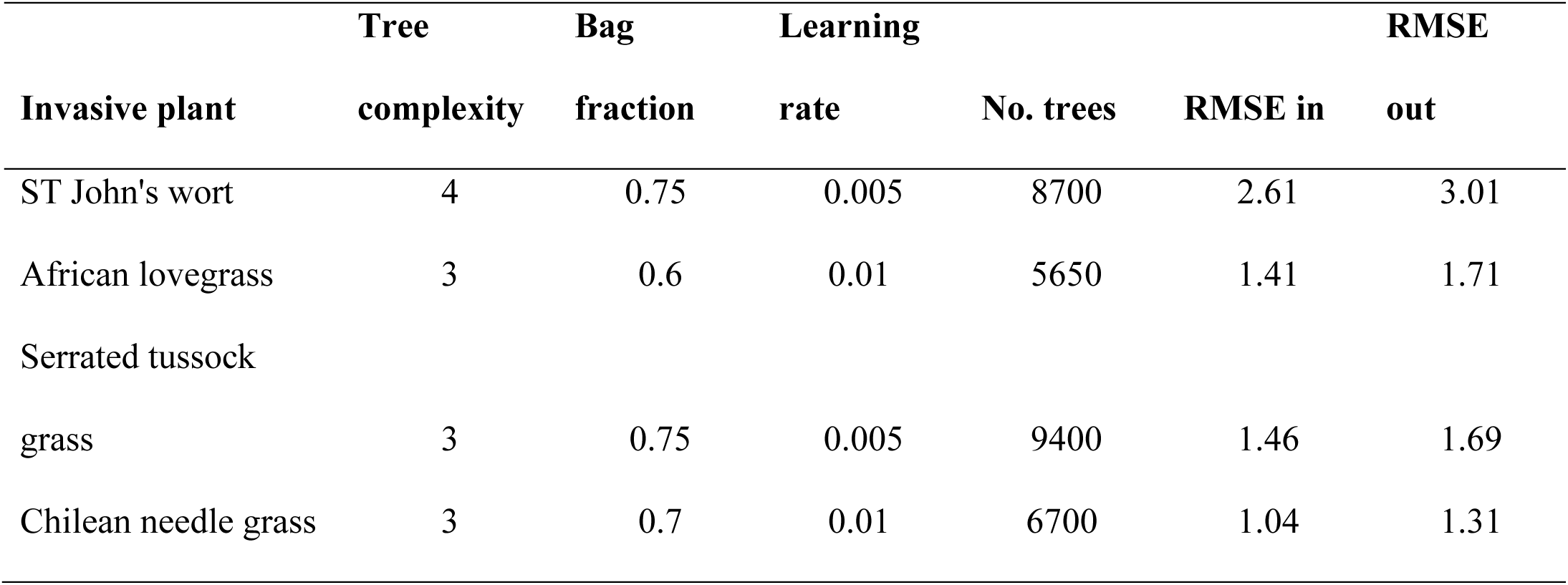
Hyperparameters and performance of the best selected model for each priority weed. For each species we tested multiple tree complexity (1 - 9), bagging fraction (0.6 – 0.75) and learning rates (0.001 – 0.03) to identify the best combination of these parameters from 144 possible combinations. Results provide for each species the root mean squared error (RMSE) for both the in-sample and out-sample performance for each species.

### Habitat suitability prediction

Across all four species, eight of the thirteen environmental covariates had comparatively higher relative influence (e.g. exceeding 5%) indicating the majority of covariates were able to capture ecologically relevant variation. The remaining covariates exhibited lower relative influence (<5%) indicating weaker effects on model predictions, although this does not necessarily imply absense of ecological relevance. Overall, we found that the multiple covariates had a relative influence that varied among species (Fig. 2). Elevation was ranked highest for both St John’s wort and Chilean needle grass, and within the top three variables for African lovegrass and serrated tussock grass (Fig. 2). Land use was an important variable for African lovegrass, but was much less influential for the other three species. Chilean needle grass showed distinct responses to several covariates, with elevation beting approximately twice as influential, fire having a stronger effect, and rainfall being less influential compared with other species. Soil process was influential for St John’s wort, but less so for the other three, and hydrogeological soil types were influential for serrated tussock grass but only marginally for the others.

**Figure 2.**
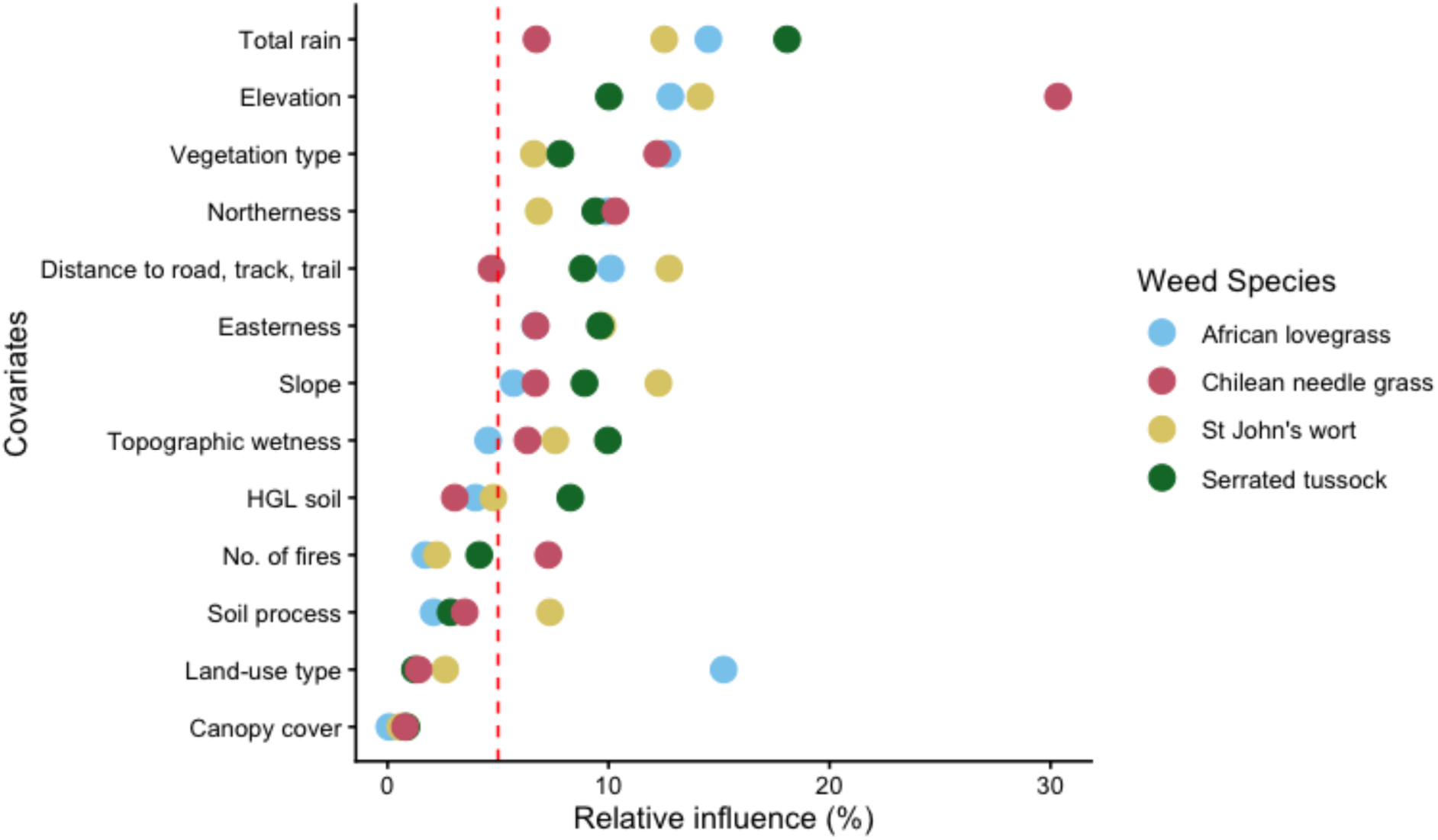
The relative influence of each environmental variable in characterising density for each invasive plant species. A higher relative influence indicates this variable was used more often in splitting trees during the model fitting process. The dashed red line denotes a low relative influence (e.g. <5%) of a predictor to improving model perfomance.

Predicted habitat suitability indices indicated considerable heterogeneity across the ACT, both within and among species (Fig. 3). African lovegrass showed the strongest associations with watercourses, with higher suitability scores particularly around the Murrumbidgee and Molonglo River catchments in the centre north. Chilean needle grass exhibited higher suitability scores in urban and peri-urban areas in the north-eastern ACT. St John’s wort displayed the most widespread distribution of higher capacity to reach higher densities particularly around roads, tracks and trails. In contrast serrated tussock grass showed higher suitability in elevated areas, and especially overlapping the Namadgi National Park to the south-west.

**Figure 3.**
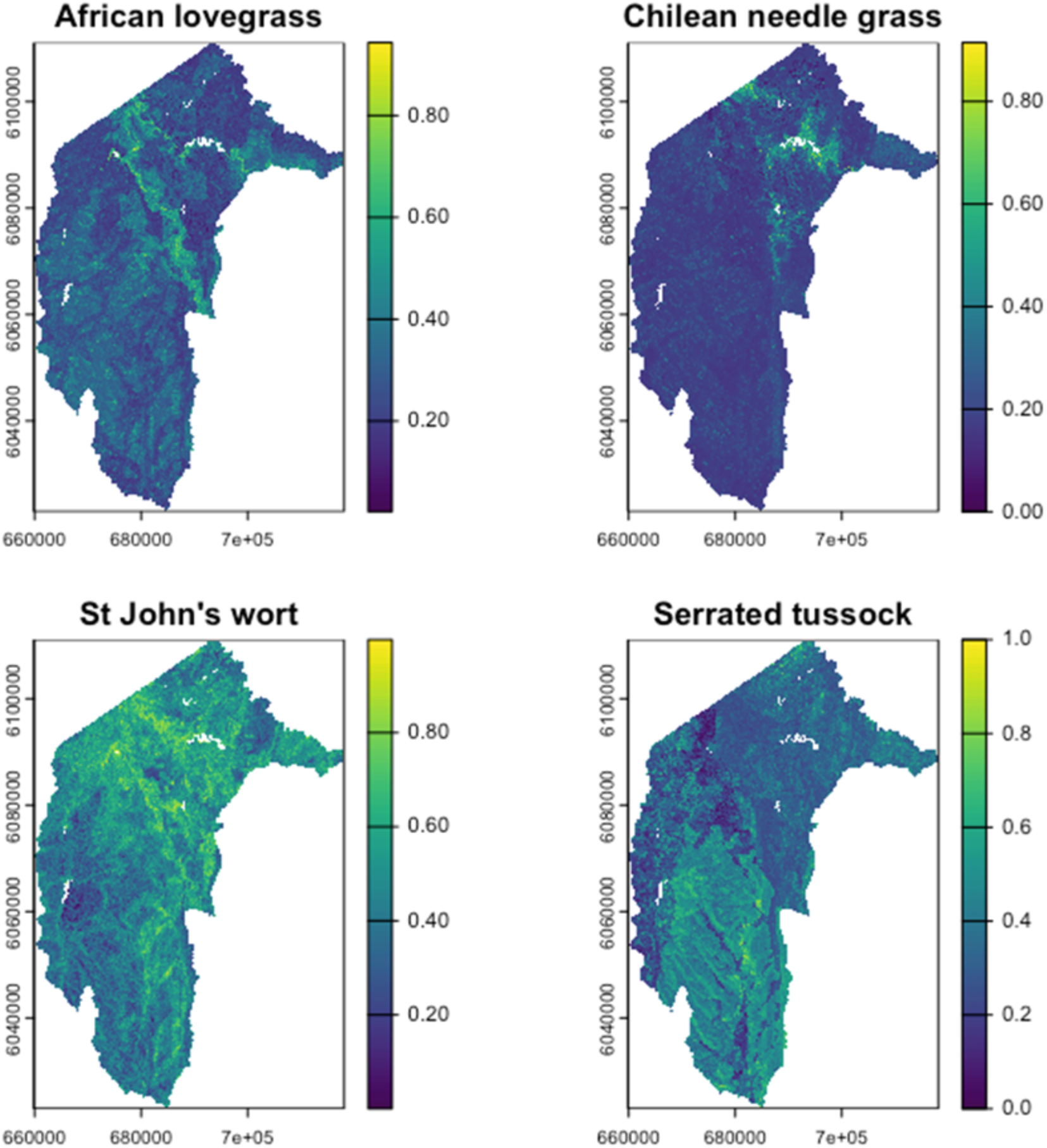
Predicted habitat suitability for four priority weeds across the Australian Capital Territory. Maps show rescaled HSI values to allow for comparisons between species. Higher values indicate areas that are predicted to be more suitable for each weed to establish and reach relatively higher densities.

### Hotspots for multi-species invasions

Overall, for all but Chilean needle grass we found more than half of the ACT was vulnerable to invasion by each weed at the mixed-density (Q1) threshold (Table 2). At the high-density (Q3) threshold, large proportions of the ACT also remained vulnerable to invasion by African lovegrass, St John’s wort and serrated tussock grass (Table 2).

**Table 2.** Comparison for the % of area within the ACT where each weed has a suitability value that exceeds a low-density (Q1) or high-density (Q3) threshold respectively.

|  | ACT % at Q1 | ACT % at Q3 |
| --- | --- | --- |
| **Invasive plant** | threshold | threshold |
| African lovegrass | 57.77 | 19.26 |
| Chilean needle grass | 16.31 | 5.44 |
| ST John's wort | 67.40 | 22.47 |
| Serrated tussock grass | 55.39 | 18.46 |

At the first threat threshold (>Q1), approximately 71% of the ACT was predicted to have overlapping suitability for two or more priority species (Fig. 4A). However, less than 4% of the ACT was predicted to be suitable for all four species. In contrast, at the higher Q3 threat threshold, only approximately 11% of the ACT was suitable for two or more priority species to reach high densities and less than 1% for all four species. These multi-species high-density invasion hotspots were scattered throughout the ACT but were most prominent along watercourses (Fig. 4B).

**Figure 4.**
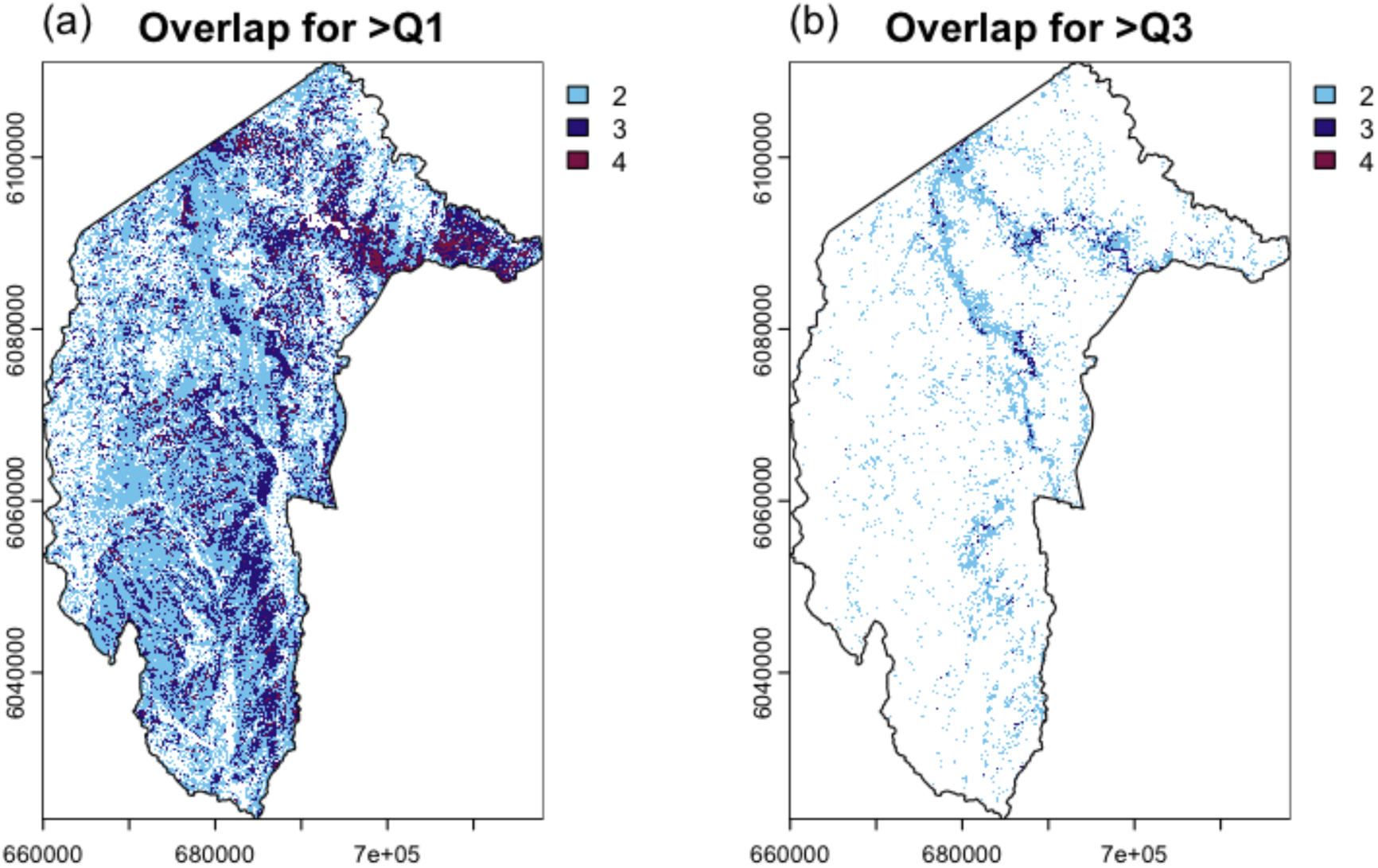
Multi-species invasion hotspots in the Australian Capital Territory. *Panel A*: shows locations where two or more out of four priority species have habitat suitability index (HSI) values exceeding the first quartile (Q1), representing areas suitable for weeds to reach low to high densities. *Panel B*: shows locations where two or more species exceeded the third quartile (Q3) threat threshold, indicating areas that are suitable for multiple species to reach only high densities.

We found, at both the Q1 and Q3 threat thresholds, that certain weeds ranked highest in predicted suitability more often than would be expected by chance (Q1 X^2^ =138091, df = 3, p =<0.001, Q3 X^2^ = 33907, df = 3, p = <0.001). At the Q1 mixed-density threshold most often raster cells were ranked highest for invasion by St John’s wort, followed by serrated tussock grass, African lovegrass and then Chilean needlegrass (Table 3). This pattern was preserved when comparing only locations for multi-species high-density threats (Q3 threshold), however St John’s wort then increased in frequency for potential locations with highest suitability for high-density invasion.

**Table 3.** Comparison for the % of locations where each weed has the highest-ranking suitability value both when exceeding the mixed low to high density threshold (Q1) and high-density threshold (Q3).

| **Invasive plant** | % Rank 1 at Q1<br>threshold | % Rank 1 at Q3<br>threshold |
| --- | --- | --- |
| African lovegrass | 7.04 | 6.83 |
| Chilean needle grass | 1.37 | 2.60 |
| St John's wort | 58.78 | 73.90 |
| Serrated tussock grass | 32.81 | 16.68 |

We found varying patterns for potential secondary invaders at the Q1 threshold for mixed density compared to the Q3 threshold for high density across the species. At the Q1 threshold, when any of the three grass species ranked highest, St John’s wort was most often ranked 2^nd^ for suitability (Fig. 5). Overall, St John’s wort displayed the widest environmental suitability, tending to overlap with the conditions favouring other species. Its suitability values as a potential secondary invader in general significantly increased in a positive direction with whichever species ranked highest at each location (mean Spearman’s rho = 0.52 + SD 0.16). At the Q3 threshold, St John’s wort remained significantly positively correlated with African lovegrass (Spearman’s rho = 0.66, p = <0.001) and Chilean needle grass (Spearman’s rho = 0.66, p = <0.001). Whereas areas where African lovegrass ranked 2^nd^ most suitable were usually where St John’s wort or serrated tussock grass were predicted to reach higher densities. However, there was not a strong signal for alignment in environmental suitability between African lovegrass and St John’s wort (Spearman’s rho = 0.13, p = <0.001), and evidence of slight divergence with serrated tussock grass (Spearman’s rho = -0.35, p = <0.00.

**Figure 5.**
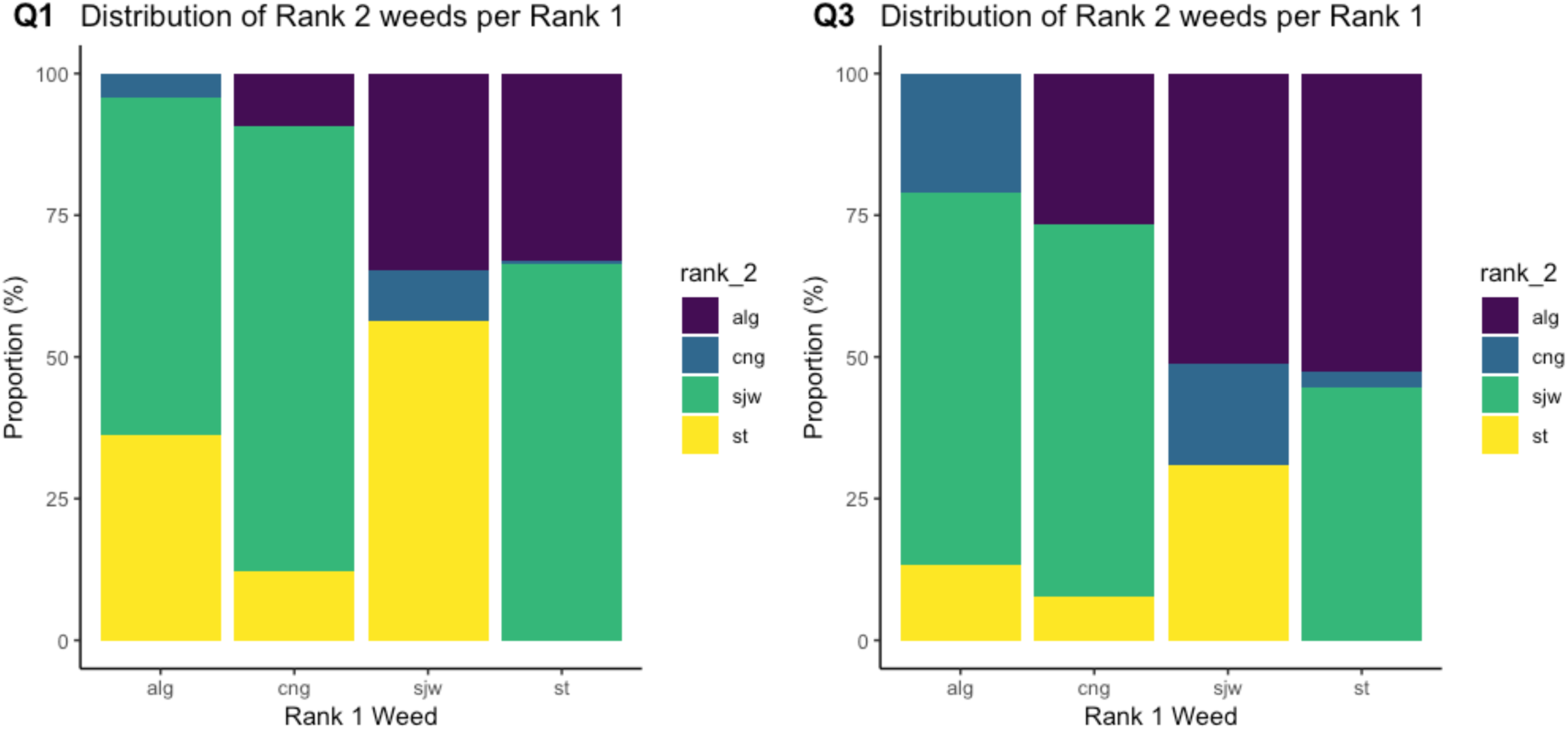
Overview of the distribution of ranking positions for each species at locations where one species ranked highest for their relative habitat suitability index, and which weed species most often ranked as second highest for their habitat suitability index at the same location.

## Discussion

Systematic monitoring of weed populations enables more strategic management by providing detailed insights into invasion threats than presence-only data. In this study, we leveraged a long-term monitoring dataset on weed density to move beyond single-species presence analyses and integrate density-based modelling with multi-species co-occurrence. This approach allowed us to not only map habitat suitability of four priority weed species but also quantify and compare potential environmental drivers influencing each species in the context of shared invasion space. This enabled us to identify invasion hotspots, areas at risk of co-occurrence or secondary invasion, and locations where targeted management interventions may improve efficiencies. These advances provide a more realistic representation of invasion dynamics at landscape scales and offer actionable guidance for integrated, spatially targeted invasive species management and planning.

### Habitat suitability predictions

Here, multiple environmental factors contributed meaningfully to habitat suitability, although their relative importance varied among species. Some variables such as rainfall, elevation, easterness, northerness and vegetation type exerted a broad influence among all species, while others such as land-use type, soils and fire varied more strongly potentially indicating species-specific associations. For example, soil process was a relatively important covariate for serrated tussock only, with the HSI map seemingly showing some association with broad soil properties. Despite being known to occur in most soil types, our result may be linked to the demonstrated sensitivity of serrated tussock to salinity (Humphries et al. 2020; Humphries and Florentine 2021). These findings also support the presence of non-linear and interactive relationships among environmental predictors for each species, consistent with the ecological complexity often associated with plant invasions (Kettenring and Adams 2011; Pyšek et al. 2012). This complexity highlights the difficulty of developing broadly generalisable management strategies.

This variability was also reflected in the spatial predictions of habitat suitability. Although sample locations and covariates were consistent across all four weed species, the models produced four distinct maps. These unique maps likely reflect variation in the ecological requirements of each species and suggests that the covariates considered represent a meaningful suite of physical, biotic and abiotic landscape properties influencing species distributions (Guisan et al. 2017a). The maps also provide geographic context for where these features combine to produce areas with the greatest suitability for each weed. In doing so, they extend managers’ perspectives beyond the local scale at which management is often implemented, supporting more holistic decision-making informed by broadscale patterns of habitat suitability.

While this variation may be largely attributed to species specific differences, it is possible other sources of variation were also contributing. Firstly, habitat suitability models operate on the assumption that the species modelled have reached equilibrium with their environment (Guisan et al. 2017c). Although our four weed species have been established in the ACT for at least 80 years, this duration may not represent equilibrium. Invasive species can often have long lag periods of 20-30 years or more (Aikio et al. 2010), meaning our estimates of suitability may be less reliable if populations are still expanding. Secondly, well-parameterised models require covariates that capture the ecological niche of a species being modelled (Guisan et al. 2017c). Although our covariates were shown to have some meaningful importance for each weed, they may not have been sampled at optimal spatial or temporal scales for these data. Some unexplained variation remained in the models, which may indicate other key covariates were missing such as seed bank survivability or dispersal. This may also have contributed to the mixed predictive performance (moderate to good) observed across species, with some models performing less strongly such as for St John’s wort. It is unclear whether the lower performance reflects inherently high ecological variability in the species, or potential biases in these data or the underrepresentation of key environmental drivers in the model. Notably, the empirical data for St John’s wort also exhibited the greatest variability and was more frequently recorded at higher densities (Supplementary Figure 1).

While these constraints should be considered when interpreting the findings, the spatially explicit patterns presented here provide useful tools to guide management on ground in an applied context. For instance, we identified areas of high habitat suitability for multiple priority species that are not yet heavily invaded but may be vulnerable to future establishment, particularly following disturbance or increased propagule pressure (e.g.

Namadgi National Park). Distance to roads tracks and trails emerged as an important driver for multiple species, with suitability often overlapping in relatively intact areas. Roadsides are known to facilitate the spread of invasive vegetation as they create disturbance through land clearing, increase light availability, and provide corridors for dispersal (Christen and Matlack 2006; Adhikari et al. 2020). Taken together, our findings suggest construction of new tracks, roads or trails within intact areas should be carefully managed to reduce invasion risk.

Preventing the negative impacts of invasive species on native ecosystems may require a high level of surveillance and our results highlight opportunities for more efficient, integrated management where resources are constrained. We acknowledge that, given the large spatial extent, perfect surveillance and management of these weeds would be impractical to achieve. However, previous research has shown that habitat suitability models can serve as useful tools for large-scale management programs to guide decisions around spatial prioritisation for surveillance and control (Richter et al. 2013). When there are resource limitations, there is a trade-off between investing in surveillance vs control. In a simulation study, Richter et al. (2013) compared management regimes based on a minimum habitat suitability threshold, which determined what areas to prioritise for surveillance vs control. Their results showed management targeting high suitability areas was most impactful in the short term, whereas including lower-suitability areas improved long-term cost-effectiveness and management success (Richter et al. 2013). Here we have provided two threat thresholds to delineate potential areas for spatially targeted management that may facilitate prioritisation of either control or surveillance of locations expected to be suitable for multiple high-threat weeds.

The usefulness of each threshold will depend on management objectives and resource availability. It is also important to note these management relevant thresholds were set arbitrarily; however, they could be refined through expert input or adjusted to better suit specific management contexts e.g. when weed density thresholds for intervention are known *a priori*.

### Hotspots for multi-species invasions

In areas where problematic invasive species have become established, a critical management priority is reducing invasion pressure from existing populations and preventing new incursions. Designing management plans for a single species across a heterogenous landscape is already a formidable challenge, integrating multiple invasive species adds further complexity yet this is the reality many managers are facing. Our results indicate that a large part of the ACT (71%) may be suitable for the establishment of multiple species, with a smaller subset (11%) potentially suitable for supporting high-density invasions. Fortunately, the worst-case scenario where habitat suitability was high for all four problematic species to reach high densities was rare (less than 1%). However, even where multiple weeds overlap at low densities, their combined effects may accumulate and lead to greater impacts. For example, Vujanović et al. (2022) found for riparian ecosystems that co-occurrence of three weed species impacted soil functioning, soil properties, and native plant diversity more than the presence of only one species. Given the overlapping distributions and potential for interactions among the four priority species examined here, coordinated management that addresses multiple species simultaneously where feasible, is likely to be more effective than single-species approaches.

Often most secondary invaders are disturbance-adapted species that opportunistically colonise newly available space (Pearson et al. 2016b). This creates challenges for management when eradication of a target weed is followed by colonisation by another high-threat species. Here we considered areas where habitat suitability overlaps to indicate secondary invasion threat among species that are already known to suppress vegetation when established at high abundance. Focusing on these priority invaders, the analysis constrains secondary invasion threat to those species that are most relevant for management, making findings directly actionable and more easily integrated into strategic planning. Our results indicate that much of the ACT was suitable for St John’s wort to reach high densities. Even in locations that were ranked as more suitable for other weeds, St John’s wort was most likely to rank second. This suggests that St John’s wort may have the broadest environmental tolerances among the four species examined. Consequently, in areas where other weeds are controlled overall St John’s wort may be the next most likely secondary coloniser. This may complicate management regimes as control methods and their costs often differ for herbs vs grasses. Biotic interactions and competitive dynamics will ultimately influence the success of secondary invaders, as outcomes depend on competitive dominance and community hierarchy (Pearson et al. 2016b; O’Loughlin and Green 2017). Nevertheless, these results provide some insight into secondary invasion threat based on relative habitat suitability and may help managers anticipate and monitor likely replacement species during weed control programs.

## Conclusions

This study advances invasion threat modelling by moving beyond single-species, presence-only approaches to demonstration of integrating density-based, multi-species habitat suitability at landscape scales. Overall, our study demonstrates the value of using weed density data to reveal patterns of invasion risk that are not detectable with conventional approaches to generate more informative habitat suitability models of long-established species requiring ongoing management. By examining four high-priority weeds simultaneously, we identified areas of potential invasion hotspots, regions vulnerable to establishment at management relevant threat thresholds, and locations where secondary invasion threat may occur. These spatial insights highlight the importance of coordinated, multi-species management approaches and emphasise the role of targeted surveillance and control in landscapes where resources are limited. The framework presented here provides a practical basis for integrating ecological modelling with adaptive management. More broadly, this work demonstrates how systematic monitoring datasets can transform invasive species management from reactive, species-by-species responses to proactive, landscape-scale prioritisation.

## Acknowledgements

This work was supported by internal funding from CSIRO as part of the APAIR portfolio. The long-term weed density monitoring was established by the Office of Nature Conservation and the Parks and Conservation Service within the ACT Government, and we extend our thanks to the many contributors for sharing these data and their efforts in collecting these valuable long term field data. We also wish to acknowledge Cameron Gallagher for his assistance and guidance with these data. This paper was improved with useful discussions with Felipe Albornoz Ramirez, Prof. Richard Duncan and XX anonymous reviewers.

## Statements & Declarations

### Funding

This research was funded through the CSIRO APaIR investment: Future Technologies for Australia’s Biosecurity with in-kind support the Australian Capital Territory Government.

## Competing interests

All authors have no competing interests to declare.

## Data and code availability

Data and code associated with this manuscript have been deposited in a public data repository and are currently embargoed but will be made accessible after manuscript acceptance.

## Authors’ contributions

All authors contributed to the study conception and design. Data preparation, and analysis were performed by KP, and WHY. The first draft of the manuscript was written by KP and all authors commented on previous versions of the manuscript. All authors read and approved the final manuscript.

## Notes

### Competing Interest Statement

The authors have declared no competing interest.

