## Supplementary files for "Modelling habitat suitability for multiple priority weed species to predict invasion hotspots for strategic management in complex landscapes"

**Biological Invasions**

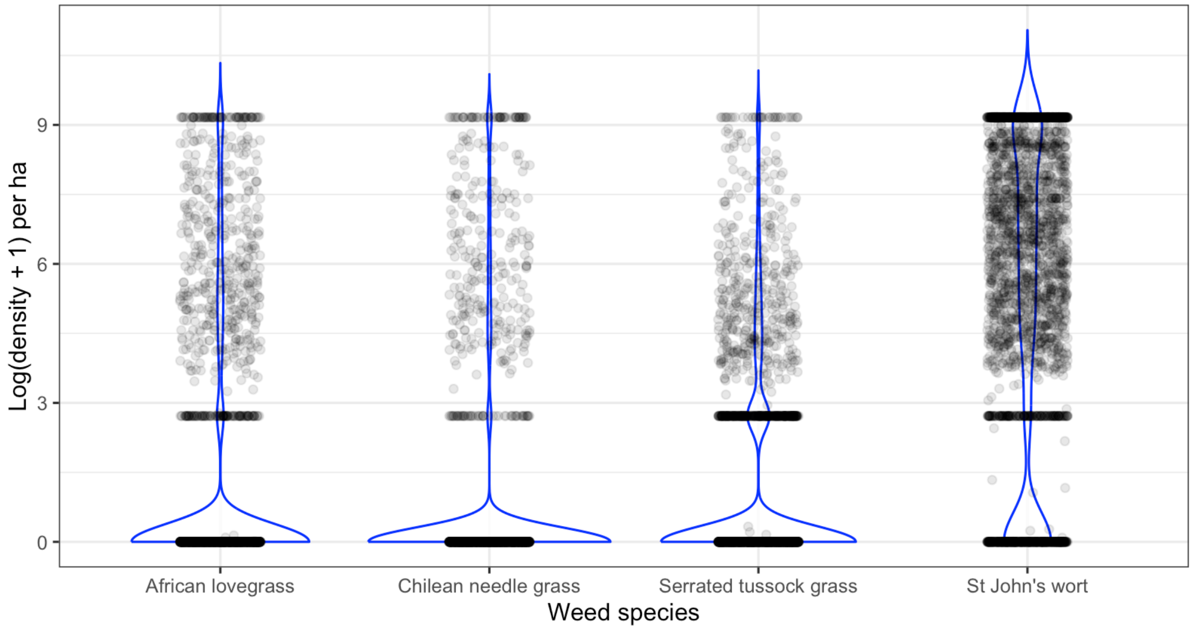


**Supplementary Figure 1**. Empirical weed density data taken from monitoring points around the Australian Capital Territory showing the upper and lower bounds in the data.

**Supplementary Table 1**. Initial modelling variables that were considered for model building. Final set of variables used for modelling (see Supplementary Table 2) were determined through a process of data driven and expert selection.

| Variable name | Unit | Description |
|---|---|---|
| soil_process | category | broad soil formation process |
| hgl_soil_ma_desc | category | hydrogeological landscape classification with geomorphic setting and dominant surface process |
| hgl_soil_d_soil_gsg | category | hydrogeological landscape classification with general soil group |
| hgl_soil_d_soil_gsg_reclass | category | simplified hydrogeological landscape classification indicating general soil group |
| hgl_soil_d_soil_asc | category | hydrogeological landscape classification with ASC |
| hgl_soil_d_soil_asc_reclass | category | simplified hydrogeological landscape classification with ASC |
| land_use_p | category | land use category |
| land_use_current_li | category | current legal/planning status of land in ACT |
| land_use_district_n | category | ACT districts |
| pct_zones_quality | code | quality of plant community type |

| | | |
|---|---|---|
| pct_zones_unstry_dom | category | dominant vegetation type in the understory of plant community type |
| pct_zones_mature_tre | category | mature trees in the plant community types |
| pct_zones_forb_diver | category | plant community type forb diversity |
| pct_value | code | plant community type value |
| zone | code | ACT zonation |
| pct_zone | code | plant community type zonation |
| veg_comm | category | vegetation community |
| veg_structure | category | vegetation structure |
| veg_formation | category | vegetation formation |
| veg_class | category | vegetation class |
| grassland_veg_com | category | grassland vegetation community |
| grassland_cons_cat | category | grassland conservation category |
| grassland_epbc_status | category | grassland EPBC status |
| grassland_act_status | category | grassland ACT status |
| grassland_category | category | grassland category |
| days_since_fire | days | number of days since fire |
| n_fires | integer | total number of fires |
| dist2road_m | meters | distance to nearest road |
| dist2trails_m | meters | distance to nearest trail |
| dist2road_track_trail | meters | distance to nearest road or trail |
| slope | degrees | slope |
| elevation | meters | elevation |
| act_slopepct3s | percent | slope |
| rain_sum_prev_30d | mm | total rainfall for previous 30 days |
| rain_mean_prev_30d | mm | mean rainfall for previous 30 days |
| rain_sd_prev_30d | mm | standard deviation of rainfall for previous 30 days |
| rain_sum_prev_90d | mm | total rainfall for previous 90 days |
| rain_mean_prev_90d | mm | mean rainfall for previous 90 days |
| rain_sd_prev_90d | mm | standard deviation of rainfall for previous 90 days |
| rain_sum_prev_365d | mm | total rainfall for previous 395 days |
| rain_mean_prev_365d | mm | mean rainfall for previous 365 days |
| rain_sd_prev_365d | mm | standard deviation of rainfall for previous 365 days |
| maxt_mean_prev_14d | degrees celcius | mean maximum temperature for previous 14 days |
| maxt_max_prev_14d | degrees celcius | maximum maximum temperature for previous 14 days |
| maxt_mean_prev_30d | degrees celcius | mean maximum temperature for previous 30 days |
| maxt_min_prev_30d | degrees celcius | maximum maximum temperature for previous 30 days |
| mint_mean_prev_14d | degrees celcius | mean minimum temperature for previous 14 days |

| | | |
|---|---|---|
| mint_min_prev_14d | degrees celcius | maximum minimum temperature for previous 14 days |
| mint_mean_prev_30d | degrees celcius | mean minimum temperature for previous 30 days |
| mint_min_prev_30d | degrees celcius | minimum minimum temperature for previous 30 days |
| vp_mean_prev_7d | hPa | mean vapor pressure for previous 7 days |
| radn_mean_prev_7d | MJ/m² | mean radiation for previous 7 days |
| evap_mean_prev_7d | mm | mean evaporation for previous 7 days |
| topo_wetness | index | topographic wetness index |
| easterness | index | the east-west orientation of a slope |
| northerness | index | the north-south orientation of a slope |
| canopy_cover | integer | A binary variable for whether the area has a foliage cover for trees over 3m or not |

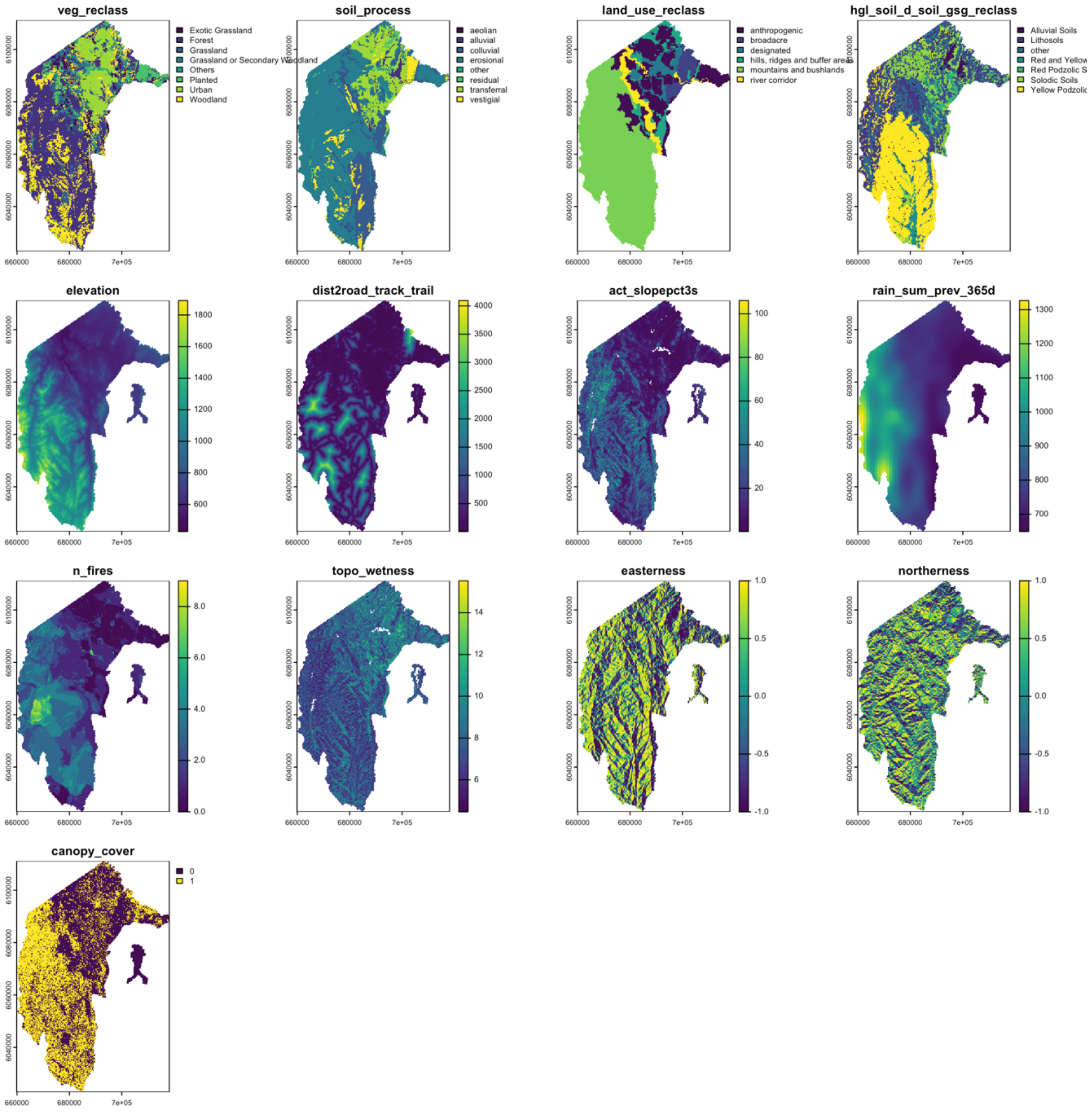


**Supplementary Figure 2.** Overview of the 13 covariates used in the development of habitat suitability maps for each of the four invasive plants. These covariates were chosen based on their representation of important environmental niche space relating to nutrient, water, sunlight and environmental disturbance factors and on their spatial and temporal alignment with the invasive plant monitoring dataset.

**Supplementary Table 2.** Details of the environmental covariate data and their respective sources that were used in the Boosted Regression Trees to create habitat suitability models for four priority weed species in the ACT. All environmental input layers were resampled to 1 hectare resolution.

| Model covariate | Description | Source | Native resolutio n |
|---|---|---|---|
| Vegetation type | Reclassified vegetation type categories including: exotic forest, exotic grassland, grassland, grassland or secondary woodland, open forest, others, planted, urban, and woodland. Modified from Vegetation 2018 mapping with an original 19 categories through expert elicitation. | ACTMapi | NA |
| Soil process | Reclassified soil processes categories including: alluvial, colluvial, erosional, other, residual, transferral and vestigial. Modified from ACT soil landscapes from 13 original categories through expert elicitation. | ACTMapi | NA |
| Land-use type | Reclassified land-use categories including: anthropogenic, broadacre, hills ridges and buffer areas, river corridor, urban open space. Modified from ACT land-use type from 24 original categories through expert elicitation | ACTMapi | NA |
| HGL soil type | Reclassified hydrogeological soil categories including: alluvial soils, lithosoils, other, red and yellow podzolic soil, red podzolic soil, solodic soil, yellow podzolic soil. Modified from hyrdological soil landscapes with 10 original categories through expert elicitation | ACTMapi | NA |
| Number of fires | Number of fires that have occurred in the ACT | Supplied by ACT Government | NA |
| Elevation | Elevation measured in meters | Supplied by ACT Government | 1m |
| Distance to road, track or trail | Distance to the nearest road, track or trail measured in meters | Supplied by ACT Government | 1m |
| Mean of annual total rainfall | Yearly total rainfall averaged over 2017 – 2023 measured in mm | NCI | 1km |
| Slope | Slope of terrain measured in percent | CSIRO DAP | 90m |
| Eastness | Topographic metric derived from a digital elevation model that measures the east-west orientation of a slope | Supplied by ACT Government | 1m |
| Northness | Topographic metric derived from a digital elevation model that measures the north-south orientation of a slope | Supplied by ACT Government | 1m |

OFFICIAL

| | | | |
|---|---|---|---|
| Topographic wetness index | Topographic Wetness Index (TWI) is calculated as log_e(specific catchment area / slope) and estimates the relative wetness within a catchment. | CSIRO DAP | 90m |
| Presence of projective foliage | A binary variable for whether the area has a foliage cover for trees over 3m or not | Supplied by ACT Government | 1m |

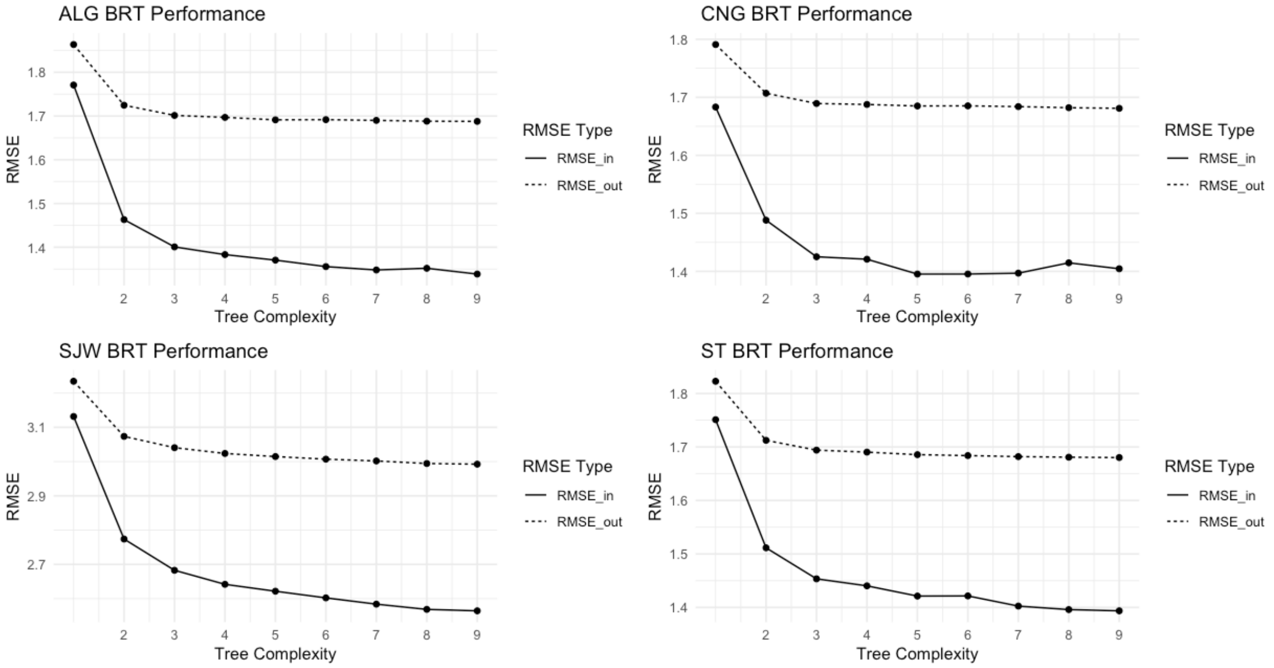


**Supplementary Figure 3**. RMSE in sample vs out sample for the chosen learning rate, and bagging fraction for each weed species from the hyper-tuning process. Showing range of tree complexity values for each weed species and the asymptote of performance. Want a small absolute RMSE and small relative gap between in-sample and out-sample to ensure performance without overfitting.

OFFICIAL

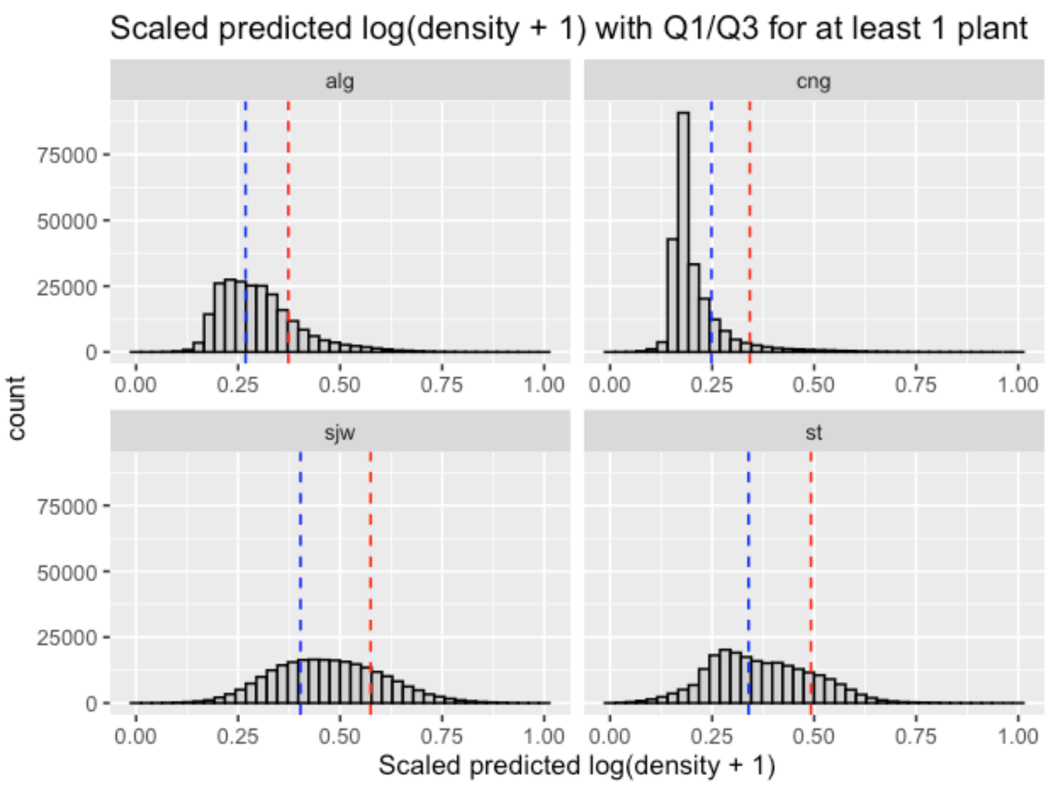


**Supplementary Figure 4.** Distribution of the predicted habitat suitability index values for each weed and the location of the two threat thresholds used to investigate multi-species hotspots. Blue dashed lines depict the threshold for exceeding the 1$^{st}$ quartile, and red dashed lines depict the threshold for exceeding the 3$^{rd}$ quartile per species.
